# Facilitation-Competition Tradeoffs Structure Microbial Niches and Nitrogen Cycling

**DOI:** 10.1101/2025.11.07.687282

**Authors:** Liang Xu, Xin Sun, Emily Zakem

**Affiliations:** Division of Biosphere Sciences and Engineering, Carnegie Institution for Science, Pasadena, CA, USA; Department of Biology, University of Pennsylvania, Philadelphia, PA, USA

## Abstract

The marine nitrogen cycle is regulated by ecological interactions among diverse microbial populations, especially in oxygen minimum zones where populations carrying out anaerobic metabolisms, mainly denitrification and anammox, drive fixed nitrogen loss. While competition for limiting resources is well studied, the combined effects of competition and facilitation, where a “feeder” population supplies a required resource to a “recipient”, remain poorly understood. Here we develop a trait-based consumer–resource framework to test how recipient populations reshape the ecological niches of their feeders and competitors. Invasion analysis shows that recipients expand either their feeder’s or the feeder’s competitor’s niche, depending on the populations’ relative competitive abilities on limiting resources. In terms of biogeochemistry, ecological outcomes result in diverse N-loss pathways; specifically, when growth is limited by both organic matter and nitrate, we observe increased nitrous oxide production. Additionally, the model suggests that anammox bacteria occupy a wider range of organic matter and nitrate supply regimes than denitrifying populations, consistent with their more frequent detection across diverse marine environments. The results link microbial interaction networks to biogeochemical fluxes relevant at global scales, and extend ecological theory to multi-resource systems with nested competitive and facilitative interactions.

## 1. Introduction

Microbial interactions regulate the transformation of energy and matter in ecosystems, from soils to oceans, and strongly influence global biogeochemical cycles (Falkowski *et al*. 2008; Bardgett & van der Putten 2014; Bertolet *et al*. 2024). In oxygen-deficient regions of the ocean, the nitrogen cycle is shaped by anaerobic microbes that transform reactive nitrogen into inert gases, driving ecosystem-scale nitrogen loss (Ward *et al*. 2009). Understanding these processes is essential for predicting how nitrogen loss will evolve, impacting primary productivity and the emission of the greenhouse gas nitrous oxide (N_2_O).

Ecological theory has provided powerful tools for studying microbial coexistence (Tilman 1982; Chesson 2000; Chase & Leibold 2003), but most models reduce complexity by focusing on competition for a single resource, treating multiple resources as substitutable, or representing facilitation as symmetric mutualism, where both species derive benefit from the interaction (Bruno *et al*. 2003; Bulleri *et al*. 2016; Orr *et al*. 2025). However, in reality, facilitation often involves intricate, asymmetric interactions rather than simple bidirectional benefits. These simplifications cannot anticipate the ecological outcomes in systems where microbes are limited by multiple, non-substitutable resources, and where interactions are competitively and facilitatively nested. How such dependencies shape both community structure and ecosystem function remains poorly understood.

Oxygen minimum zones (OMZs) provide a natural laboratory for advancing microbial interaction theory. In these anoxic waters, two main functional groups mediate most fixed nitrogen loss: heterotrophic denitrifying microorganisms (hereafter, “denitrifiers”) and anaerobic ammonium-oxidizing (anammox) bacteria (Lam & Kuypers 2011). Denitrifiers use nitrate 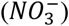 or nitrite 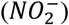 as electron acceptors to oxidize organic matter, producing a stepwise cascade of intermediates—from nitrite and nitric oxide to nitrous oxide (*N*_2_*N*) and ultimately dinitrogen gas (*N*_2_). They are phylogenetically and functionally diverse, with different lineages performing different subsets of these steps (Sun & Ward 2021; Zhang *et al*. 2023). Some produce only intermediates (e.g., *N*_2_*O*), while others carry reactions to completion (Fig. 1). In contrast, anammox bacteria are phylogenetically conserved autotrophs that fix carbon by oxidizing ammonium 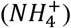 with nitrite, producing *N*_2_ as their sole product. These functional groups are metabolically interconnected, with nitrite produced by some denitrifiers (the feeders) providing a key substrate for both other denitrifiers and anammox bacteria (the recipients). Consequently, shifts in the abundance or traits of one group, set by external forcings such as nutrient supply, can cascade through the community, restructuring coexistence and altering the magnitude and form of nitrogen loss.

**Figure 1.**
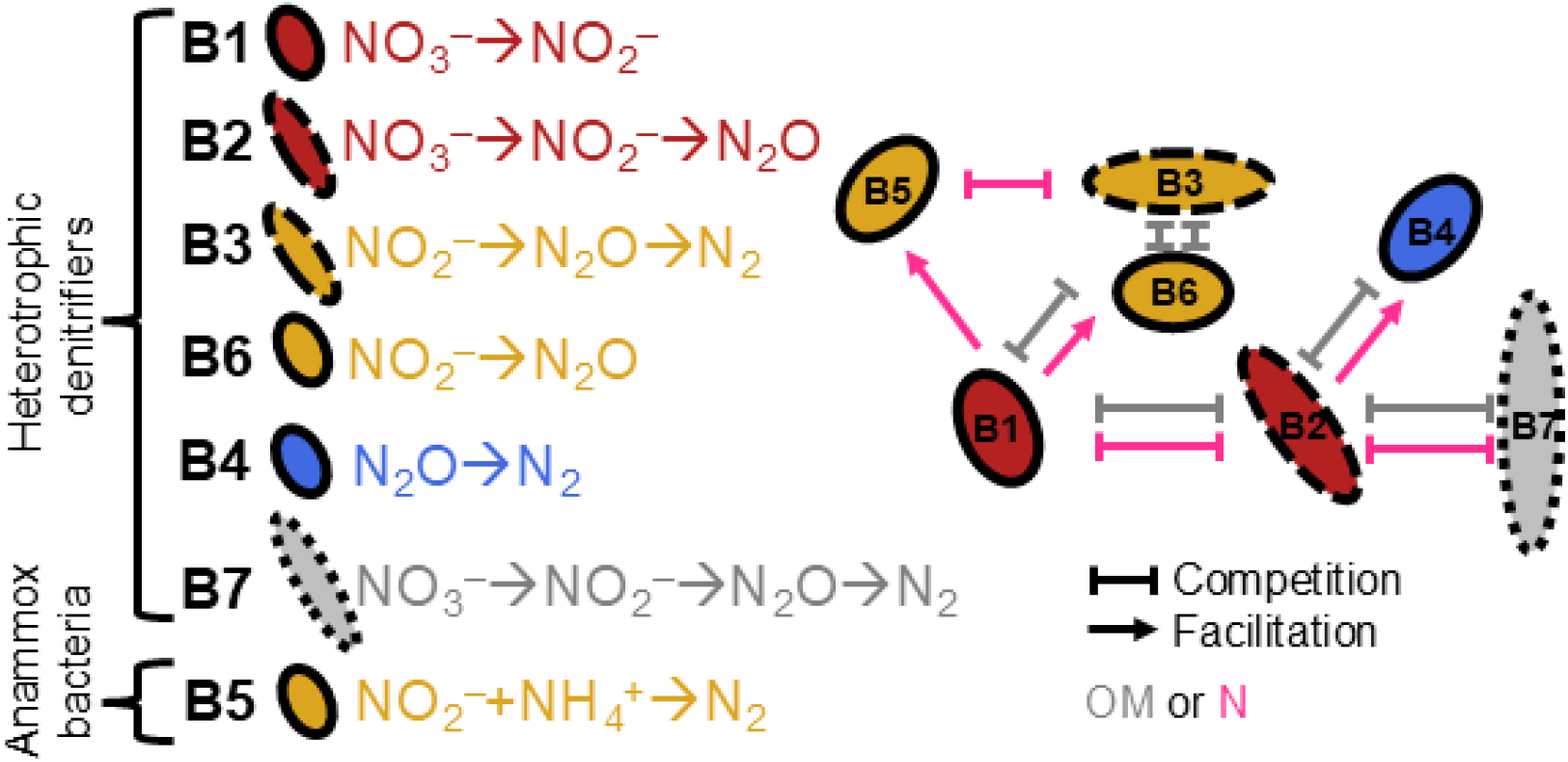
Microbial functional groups represented in the model and a schematic summary of their interactions.

In this study, we analyze a mechanistic consumer–resource model that links microbial traits to the supply and transformation of organic matter and nitrate. Microbial populations carry out diverse N-cycling metabolisms that are quantitatively distinguished from one another using the underlying redox chemical reaction (Sun et al. 2024). Using invasion analysis and graphical theory, we ask how the presence of recipient populations modifies the coexistence space of their feeders and competitors, and how these changes propagate to ecosystem-scale nitrogen loss. Our results show that recipients can either contract or expand a feeder’s niche depending on their resource use, and that the relative supply of organic matter and nitrate governs whether nitrogen is lost primarily as dinitrogen or as nitrous oxide. Our analysis extends beyond that of Sun et al. 2024 to consider the two-dimensionality of the gradients in organic matter and nitrate supply, which allows us to identify a region of co-limitation with a qualitatively different ecological outcome where only nitrous oxide is produced. We also more comprehensively consider the role of anammox as both a competitor and a recipient. By linking microbial ecology with biogeochemistry, our framework provides new insights into the mechanisms that regulate nitrogen cycling in the ocean. Such theoretical advances are critical for interpreting observed patterns of nitrogen loss and understanding how they change over time.

## 2. Methods

### 2.1. Functional Groups

We consider seven microbial functional groups that represent the major anaerobic nitrogen-cycling pathways in oxygen-deficient marine systems (Sun & Ward 2021; Zhang *et al*. 2023). These include the stepwise denitrifiers, anammox bacteria, and the complete denitrifier. Each group is defined by its essential resource requirements—typically organic matter (OM) as an electron donor and nitrate or nitrite as acceptors—and by the metabolic byproducts they release. These byproducts frequently serve as substrates for other groups, establishing a network of competitive and facilitative interactions. Presenting the functional types first provides the biological context for how resources flow among populations and why dependencies arise among them (Fig. 1).

1. 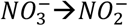 denitrifiers (Group 1, *B*_1_): Heterotrophic denitrifiers that use OM and nitrate 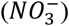 to produce nitrite 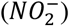.
2. 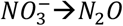 denitrifiers (Group 2, *B*_2_): Heterotrophs that use OM and 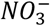 to produce nitrous oxide (N_2_O).
3. 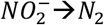 denitrifiers (Group 3, *B*_3_): Heterotrophs requiring OM and 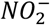, producing dinitrogen gas (*N*_2_).
4. *N*_2_*O*→*N*_2_ denitrifiers (Group 4, *B*_4_): Heterotrophs requiring OM and *N*_2_*O*, producing *N*_2_.
5. Anammox bacteria (Group 5, *B*_5_): Autotrophs that oxidize ammonium (NH_4_^+^) with nitrite 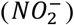, using the released energy to fix inorganic carbon, and producing *N*_2_ as their sole product.
6. 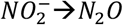 denitrifiers (Group 6, *B*_6_): Heterotrophs requiring OM and 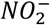, producing *N*_2_*O*.
7. Complete denitrifiers (Group 7, *B*_7_): Heterotrophs that reduce nitrate 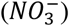 all the way to *N*_2_, using OM as the electron donor.

### 2.2. Model Framework

To formalize these interactions, we implement a virtual chemostat–based consumer–resource model that tracks both microbial biomasses and the concentrations of OM, nitrate, nitrite, nitrous oxide, and ammonium. Growth follows Liebig’s Law of the Minimum, reflecting the requirement that each functional group depends on two essential and non-substitutable resources. The model links the physiological traits of each group—maximum uptake rates, affinities, and biomass yields—to their realized growth rates and subsistence concentrations. This structure enables direct comparison of competitive abilities and allows facilitative links to emerge naturally from the excretion of metabolic byproducts defined in Section 2.1.

The dynamics are generally given by:

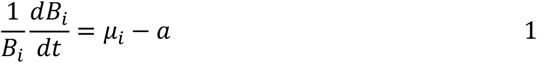

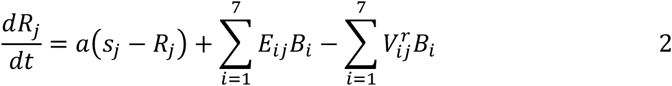

where *B*_*i*_ is the biomass of group *i*, and *a* is the dilution rate. To compute growth and uptake under Liebig’s Law, we proceed in two steps. The maximum specific uptake rate for each functional group *i* on resource *j*, 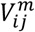, is specified a priori from the trait parameterization (Table 1). For each essential resource, we first calculate the potential growth rate it can support:

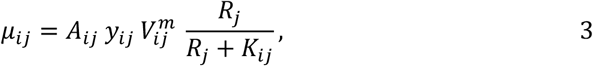

where *y*_*ij*_ is the biomass yield, *K*_*ij*_ is the half-saturation constant, and *R*_*j*_ is the resource concentration. *A*_*ij*_ indicates whether group *i* uses resource *j*, given by

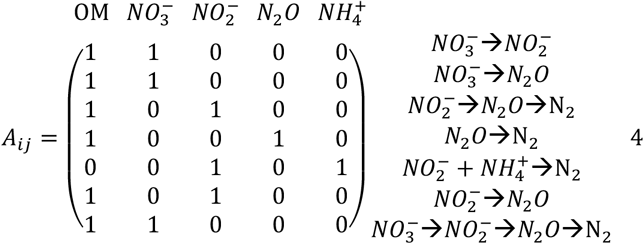

**Table 1:** Physiological trait values are identical to those used in Sun *et al*. (2024).

| Microbes | Substrate | Max Uptake rate<br>$V_{ij}^m$ (MOL N $d^{-1}$) | Yield $y_{ij}$ | Half-saturation<br>rate $K_{ij}$ ($\mu M$ N) | $R_{ij}^*$ |
| --- | --- | --- | --- | --- | --- |
| $B_1: NO_3^- \rightarrow NO_2^-$ | $R_1: OM$ | 1.377 | 0.15 | 0.1 | 0.024 |
| $B_1: NO_3^- \rightarrow NO_2^-$ | $R_2: NO_3^-$ | 50.8 | 0.011 | 4 | 0.295 |
| $B_2: NO_3^- \rightarrow N_2O$ | $R_1: OM$ | 1.377 | 0.148 | 0.1 | 0.0245 |
| $B_2: NO_3^- \rightarrow N_2O$ | $R_2: NO_3^-$ | 50.8 | 0.023 | 4 | 0.144 |
| $B_3: NO_2^- \rightarrow N_2$ | $R_1: OM$ | 1.377 | 0.197 | 0.1 | 0.0172 |
| $B_3: NO_2^- \rightarrow N_2$ | $R_3: NO_2^-$ | 50.8 | 0.024 | 4 | 0.138 |
| $B_4: N_2O \rightarrow N_2$ | $R_1: OM$ | 1.377 | 0.298 | 0.1 | 0.0108 |
| $B_4: N_2O \rightarrow N_2$ | $R_4: N_2O$ | 50.8 | 0.013 | 0.6 | 0.039 |
| $B_5: ANAMMOX$ | $R_5: NH_4^+$ | 50.8 | 0.013 | 0.45 | 0.028 |
| $B_5: ANAMMOX$ | $R_3: NO_2^-$ | 50.8 | 0.011 | 0.45 | 0.034 |
| $B_6: NO_2^- \rightarrow N_2O$ | $R_1: OM$ | 1.377 | 0.201 | 0.1 | 0.0169 |
| $B_6: NO_2^- \rightarrow N_2O$ | $R_3: NO_2^-$ | 50.8 | 0.016 | 4 | 0.2066 |
| $B_7: NO_3^- \rightarrow N_2$ | $R_1: OM$ | 1.377 | 0.138 | 0.1 | 0.0268 |
| $B_7: NO_3^- \rightarrow N_2$ | $R_2: NO_3^-$ | 50.8 | 0.026 | 4 | 0.124 |

In classic resource competition theory (Tilman 1982), the subsistence concentration 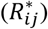 for each resource, defined as the minimum concentration at which the growth rate balances mortality, implies competitive ability to that resource:

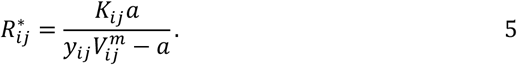

Lower 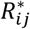 values indicate stronger competitive ability under the limitation of resource *j* (Table 1). The realized growth rate is then given by Liebig’s Law as

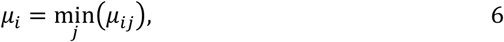

corresponding to the most limiting essential resource.

Because the realized growth rate cannot exceed the rate supported by the limiting resource, uptake of all essential nutrients must be adjusted accordingly. The realized uptake rate on resource *j* is therefore

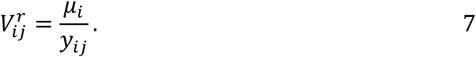

This formulation ensures that 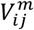 defines the potential maximum uptake, while the realized uptake 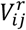 remains consistent with the growth rate imposed by the limiting resource, as required in a Liebig-type consumer–resource model.

The model tracks the concentrations of OM, nitrate 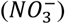, nitrite 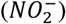, nitrous oxide (*N*_2_*O*), and ammonium 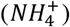, denoted by *R*_1_, ⋯, *R*_5_ respectively. External supplies (*s*_1_ and *s*_2_) are provided for OM and 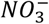. The excreted compounds of microbial metabolism are defined in an excretion matrix *E*_*ij*_, given by

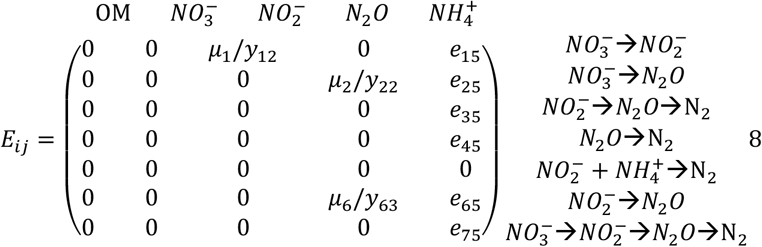

For example, 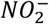 is produced by 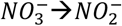 denitrifiers, and *N*_2_*O* is produced by 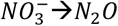 and 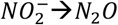 denitrifiers. Note that 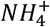 arises internally as a byproduct of OM remineralization by heterotrophs, determined by the C:N of the OM and the electron-balanced budget for each metabolic functional type (Sun *et al*. 2024). Thus, we assume that 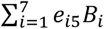 in the range of 70–86% of *s*_1_ as a supply rate of 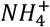.

Trait values were derived following the framework of Sun et al. (2024), which estimates the biomass yields from underlying redox chemistry and energetic constraints (Rittmann & McCarty 2012), and shows consistency with observations (Table 1).

### 2.3. Simulation Setup and Analysis

We focused on ecological equilibria—specifically, microbial survival, exclusion, and community composition—under a wide range of nutrient supply conditions. To explore how nutrient regimes shape coexistence, we systematically varied the supply rates of OM and 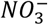 across ecologically relevant ranges (Supplementary Material).

Invasion analysis was used to determine whether a group could increase from low abundance in a given resource environment. Graphical analyses of zero net growth isoclines (ZNGIs) and consumption vectors were employed to visualize competitive outcomes and facilitative interactions among groups (Chase & Leibold 2003; Xu *et al*. 2025). The consumption vector characterizes the relative uptake of essential resources and delineates the regions of coexistence and dominance among groups (Tilman 1982; Chase & Leibold 2003; Xu *et al*. 2025). In our framework, it is defined as the ratio of yields on each of the two essential substrates for each functional type. For example, for the 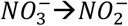 denitrifiers that utilize OM and 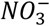 as essential substrates, the consumption vector is given by (see derivation of Eqn. S20 in the Supplementary Material):

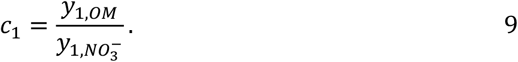

We then evaluate how the consumption vectors change when recipients from different feeder groups are introduced, to examine the resulting shifts in coexistence patterns.

We also examine how microbial interactions influence ecosystem-scale nitrogen cycling by determining the form of nitrogen loss at steady state. Nitrogen is considered lost when converted to gaseous products (*N*_2_ or *N*_2_*O*). Across most of the 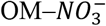 supply domain, *N*_2_*O* is produced and consumed internally at comparable rates, making it a transient intermediate rather than a true loss pathway. However, this cycling still identifies regions with high potential for *N*_2_*O* emission: if an external process— such as physical mixing or transport—were to decouple *N*_2_*O* production from its consumption, these regions could generate net *N*_2_*O* release. By mapping N-loss outcomes across the two-dimensional supply space of OM and nitrate, we identified threshold regimes, coexistence regions, and the dominant pathways under which either *N*_2_ or *N*_2_*O* (or a combination) is favored, including a distinct co-limitation zone where simultaneous OM and nitrate limitation results in net *N*_2_*O* accumulation.

## 3. Results and Discussion

We analyze the resulting complex interactions among the seven microbial functional types by incrementally constructing the system from simpler subsystems, starting with only two microbial types, 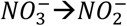 denitrifiers (*B*_1_) and 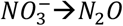 denitrifiers (*B*_2_), as all other types depend on production from one of these two types (except for a third 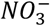 consumer, the complete denitrifier (*B*_7_), discussed below). This stepwise approach allows us to disentangle the quantitative principles that govern microbial coexistence, exclusion, and resource partitioning. Furthermore, simplified microbial consortia are central to engineered ecosystems, such as wastewater treatment facilities, where specific functional groups—particularly denitrifiers and anammox bacteria—are selectively enriched to maximize nitrogen removal efficiency (Rittmann & McCarty 2012; Ali *et al*. 2020), and so understanding the interactions of these simplified consortia could be useful for these contexts (Baeten *et al*. 2019).

### 3.1. Baseline System: Two Nitrate-reducing Competitors

We begin by analyzing the baseline system, which includes two heterotrophic denitrifiers competing for OM and nitrate 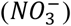: the 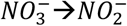 denitrifier (*B*_1_) reducing 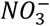 to nitrite 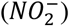 and the 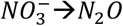 denitrifier (*B*_2_) reducing 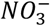 directly to nitrous oxide (*N*_2_*O*). Trait-based parameterization reveals a trade-off in their resource requirements (Sun *et al*. 2024): the 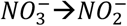 reducer has a lower subsistence requirement and a better competitor for OM 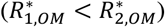 but a higher requirement and a weaker competitor for 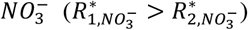 compared to the 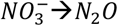 reducer (Table 1). Their consumption vectors 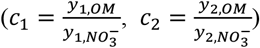 then form a coexistence region (Fig. 2a) (Tilman 1982; Chase & Leibold 2003; Xu *et al*. 2025). As a result, the OM: 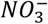 supply ratio determines competitive outcomes. When the OM: 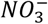 supply ratio is small, the 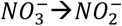 reducer dominates; while the OM: 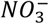 supply ratio is large, the 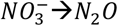 reducer prevails; and at intermediate supply ratios, both groups coexist (Fig. 2a).

**Figure 2.**
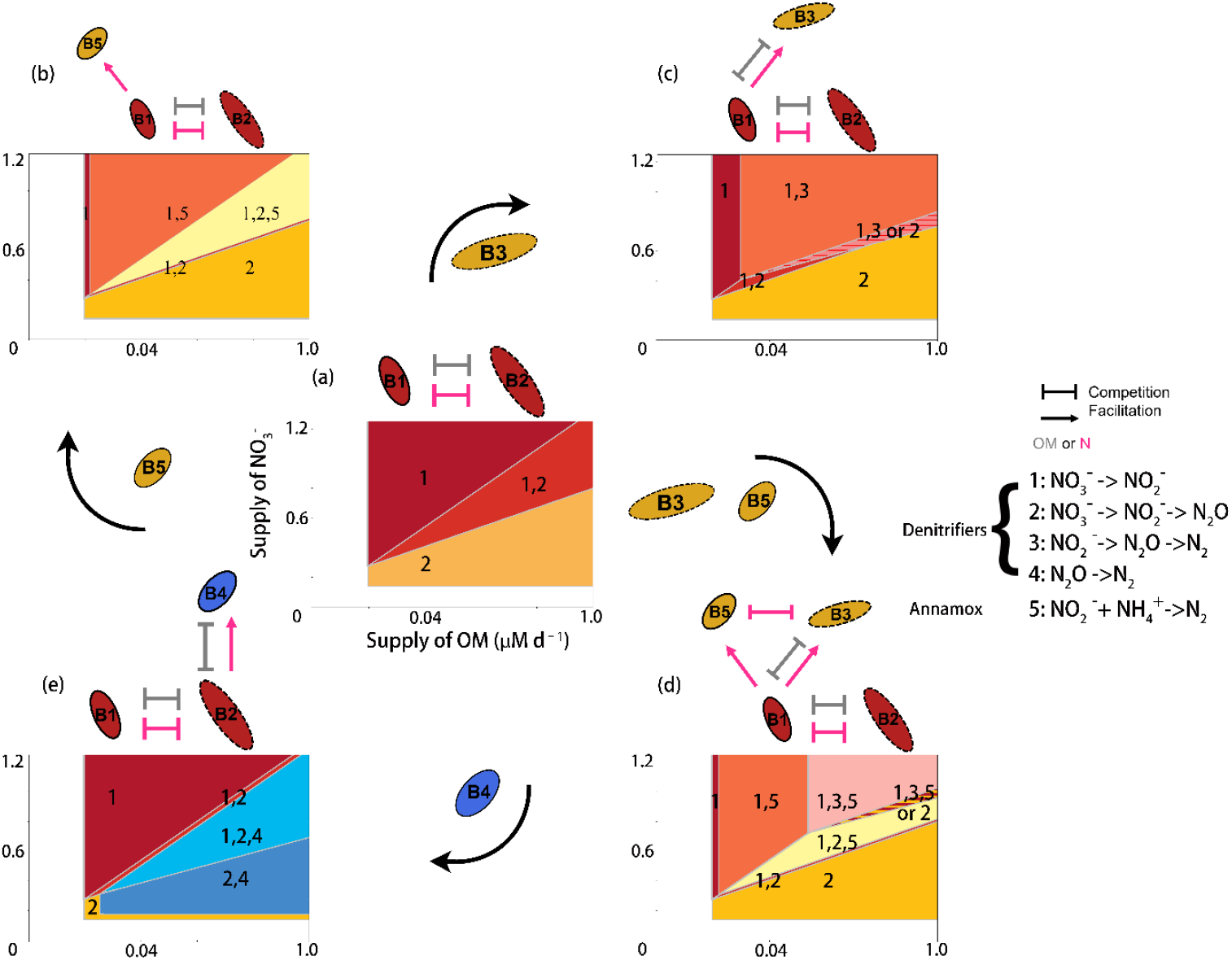
Competitive outcomes as a function of organic matter (OM) and 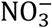 supply rates across systems. Outcomes are shown in the 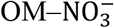 supply space for the baseline system (a, center panel) and for alternative subsystems (b–e, surrounding panels).

In this subsystem, because only the 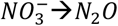 reducer directly produces *N*_2_*O*, nitrogen loss occurrs only when OM supply is sufficiently high to sustain the 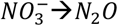 reducer’s persistence (Fig. 3a).

**Figure 3.**
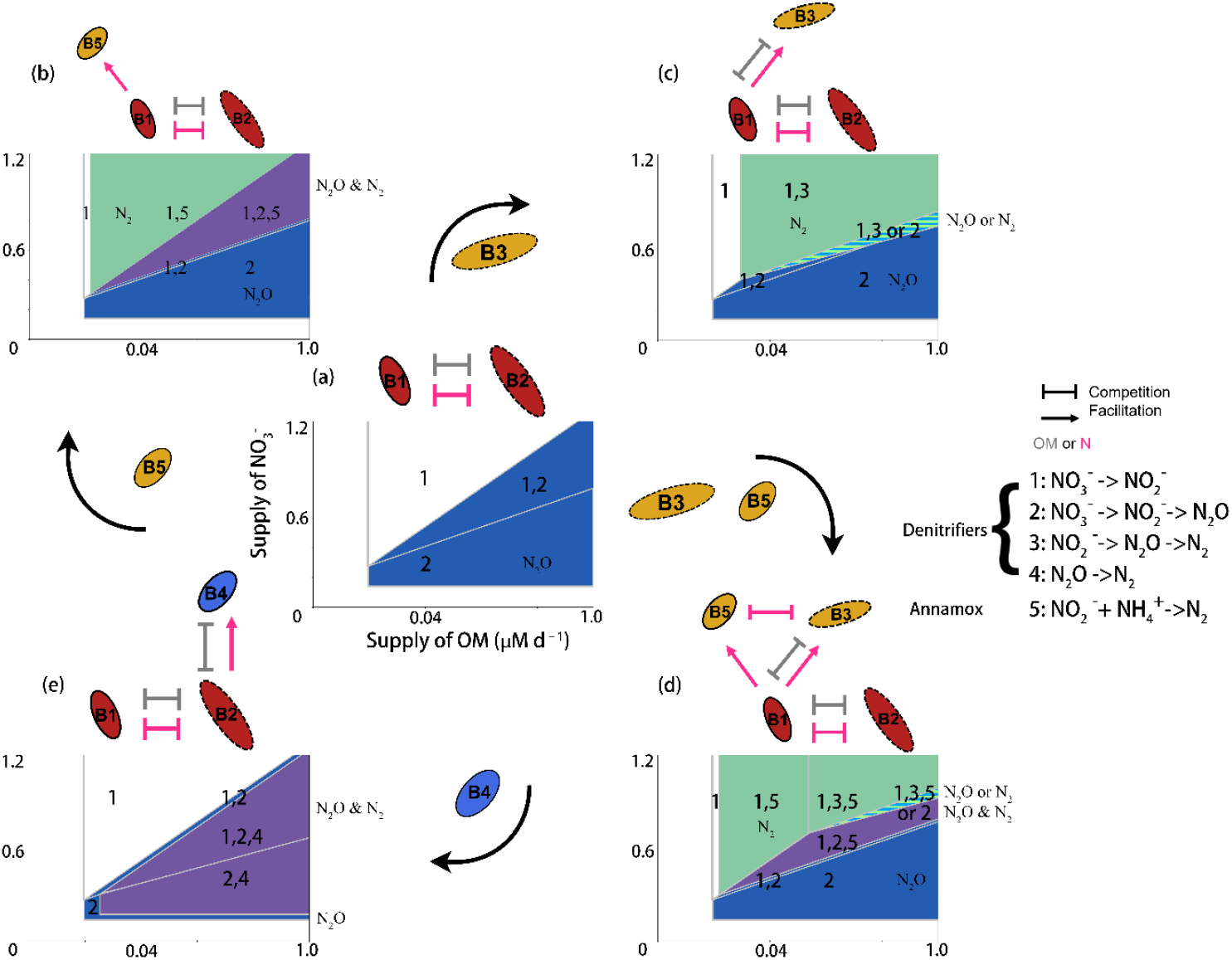
Nitrogen loss pathways as a function of organic matter (OM) and 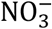 supply rates. Pathways are mapped in the 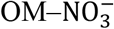 supply space for the baseline system (a, center panel) and for the corresponding subsystems (b–e, surrounding panels).

A third denitrifier—the complete denitrifier, *B*_7_—is a direct competitor with *B*_1_ and *B*_2_, using both OM and 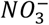. However, it has a competitive advantage in utilizing 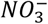 (a lower 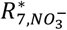), allowing it to outcompete *B*_1_ and *B*_2_ under more 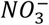 –limited conditions (Sun *et al*. 2024). Consequently, *B*_7_ dominates the community when 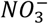 availability is low (Fig. 4a). Because its coexistence region lies largely outside the primary parameter space of the other groups, we do not further analyze this denitrifier in detail but include it in the full system for completeness.

**Figure 4.**
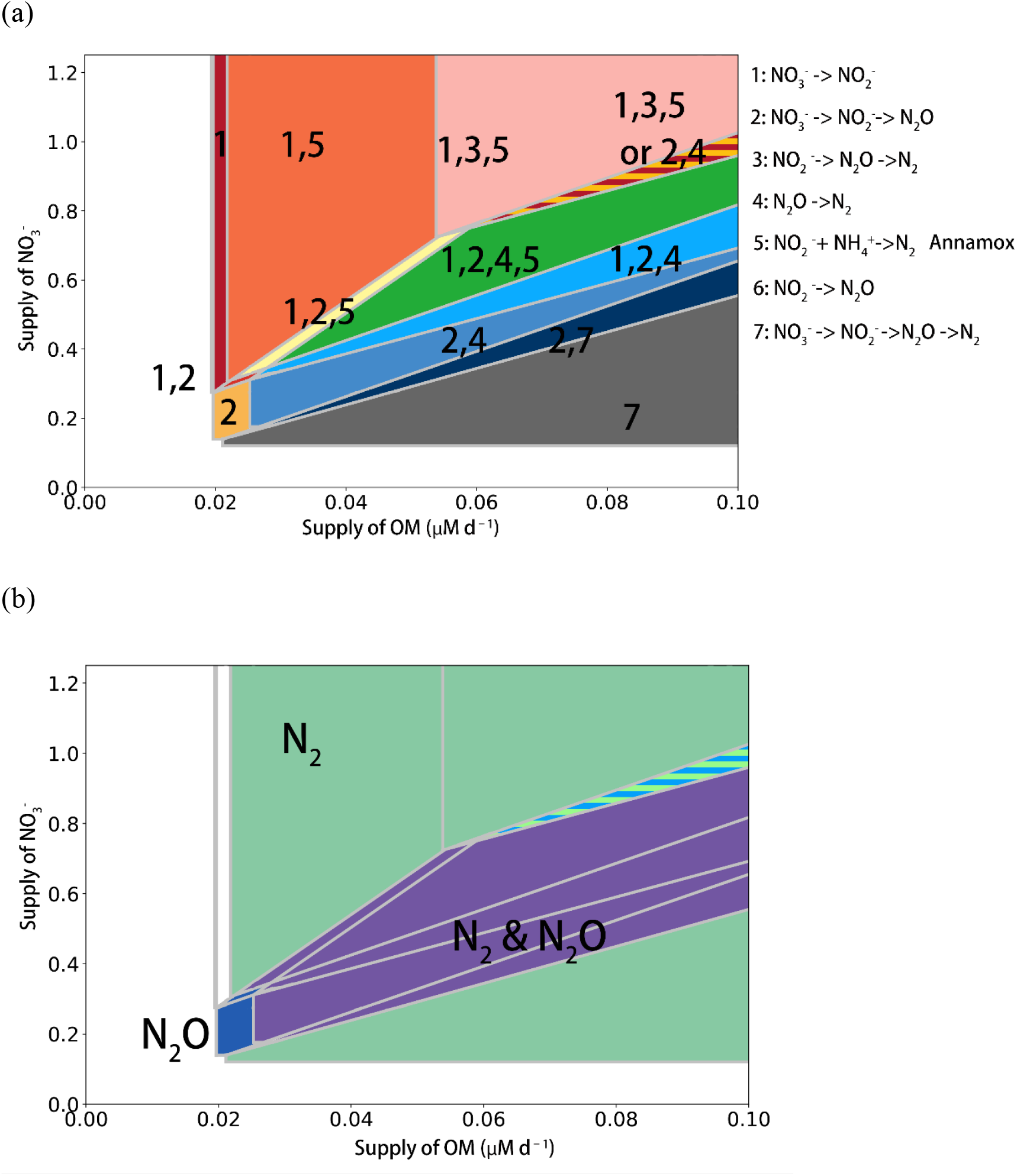
Graphical analysis of functional group coexistence (a) and nitrogen loss pathways (b) in the full system, together with the corresponding N-loss area. Nitrogen loss pathways denote the chemical form in which nitrogen is removed—either as N_2_ or N_2_O. Regions showing both N_2_ and N_2_O correspond to steady states in which N_2_O production is balanced by its consumption; in natural environments (e.g., the ocean), such conditions nonetheless create the potential for N_2_O emission.

### 3.2. Recipient Groups Restructure Competitive Outcomes

We next examine how introducing recipient functional groups alters these baseline dynamics. Recipients require substrates produced by feeders, creating facilitative links, but often also compete with feeders or feeders’ competitors for OM. These dual roles restructure coexistence outcomes in qualitatively different ways depending on resource overlap.

#### Adding Anammox: Neutral Recipient, Expanded Nitrogen Loss

We first introduce anammox bacteria, which rely on nitrite produced by the 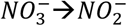 denitrifier and on ammonium derived from OM remineralization (Eqn. 4).

Anammox bacteria can be theoretically limited by one or both of their essential substrates, nitrite and ammonium. When anammox is limited by nitrite, which is produced by the 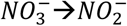 denitrifier (*B*_1_), its biomass scales with the density of *B*_1_, with a minimum requirement for the density of *B*_1_ to produce enough nitrite to sustain anammox. A general criteria for the minimum equilibrium density of a feeder type required to sustain a recipient type is given by (see the supplementary material for derivation, Eqn. S15):

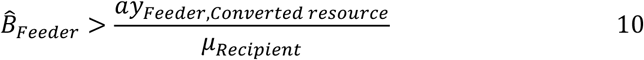

Thus, the equilibrium density of *B*_1_ required for anammox to invade is 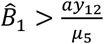 while *y*_12_ denotes the yield of the feeder *B*_1_ by synthesizing nitrate (*R*_2_), which is converted to nitrite (*R*_3_). *μ*_5_ is the realized growth rate of anammox (*B*_5_) when limited by nitrite (*R*_3_) produced by the feeder type. Eqn. 10 shows that with a constant dilution rate, the requirement of the biomass of the feeder type increases with the yield of the feeder on nitrate as more *N* is absorbed in *B*_1_ biomass and decreases with the growth rate of anammox on nitriteanammox. This is reflected in the graphic analysis as the ZNGI of *B*_5_ being above the ZNGI of *B*_1_ (Fig. 2b), showing that anammox can only persist when the 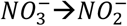 denitrifier (*B*_1_) establishes a sufficient biomass.

In contrast, if nitrite is sufficiently supplied, anammox can be limited by ammonium, supplied by the remineralization from OM from the heterotrophs. In either case, the dynamics of anammox are decoupled from the direct competition between nitrate reducers. Thus, anammox does not alter the competitive boundaries between the two nitrate reducers.

The presence of anammox expands the region of nitrogen loss by introducing a pathway for *N*_2_ production in conditions where the baseline system conserves nitrogen (Fig. 3b). The N-loss via *N*_2_ then happens in the region where anammox can persist. This case demonstrates that recipients can broaden the functional consequences of microbial communities without altering coexistence boundaries among their feeders and the feeders’ competitor.

#### Adding 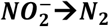 Denitrifiers: Priority Effects via Intensified Competition

Introducing the 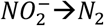 denitrifiers (*B*_3_) into the baseline system produces a distinct outcome. Similar to anammox, these denitrifiers depend on nitrite supplied by the 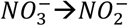 reducer. Accordingly, the establishment of *B*_3_ requires a minimum threshold density of *B*_1_, given by 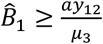 (Eqn. 10). Unlike anammox, however, *B*_3_ also consumes OM and exhibit the lowest subsistence requirement for OM among the three groups 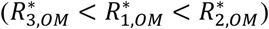 (Table 1). Consequently, the 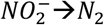 denitrifiers must remain limited by nitrite when coexisting with the 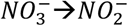 reducer; otherwise, they would exclude the latter under OM-limited conditions when 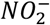 is sufficiently supplied externally.

Once established, the feeder–recipient pair forms a tightly coupled consortium that depletes OM more strongly than the 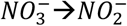 reducer alone. This is reflected in the consortium’s consumption vector (Eqn. S19)

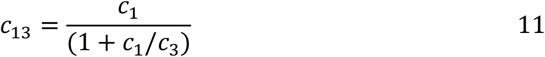

where 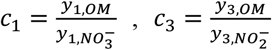 and *c*_13_ is the consortium’s consumption vector. Eqn. 11 shows that *c*_13_ is smaller than *c*_1_, indicating a greater relative consumption of OM compared to the 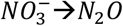 denitrifiers. Consequently, the consortium exerts a stronger suppressive effect on 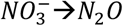 denitrifiers under OM-limited conditions, thereby narrowing their niche (Fig. 2ac).

Furthermore, the consortium’s stronger resource consumption on OM can generate priority effects in cases where 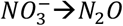 and 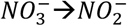 coexist in the baseline system. In this case, the outcome depends on the order of establishment (Fig. 2c and Fig. S1, Eqn. S19-S20) (Chase & Leibold 2003; Fukami 2015; Xu *et al*. 2025). If the feeder–recipient pair becomes established first, they prevent invasion by the 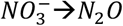 reducer; if the 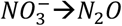 reducer arrives first, it can exclude the pair. Therefore, the niche space of the 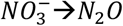 denitrifiers is potentially narrowed (Fig. 2ac).

From a biogeochemical perspective, the presence of the 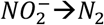 recipient expands the region of nitrogen loss by enabling *N*_2_ production under OM-sufficient conditions, similar to the subsystem containing anammox (Fig. 3ac). In regions governed by priority effects, the dominant form of nitrogen loss depends on the community’s assembly history—yielding either *N*_2_ from the feeder–dependent pair or *N*_2_*O* from the 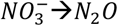 denitrifiers.

#### Interactions Among the Two Recipients of the 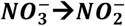 Reducer

When both recipients of the 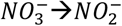 reducer—anammox and 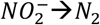 denitrifiers—were present, their coexistence outcomes depend on ammonium availability. Our study parameterizes anammox as a more efficient nitrite competitor than 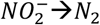 denitrifiers (Table 1), characterized by a lower minimum subsistence concentration 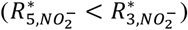. If not so (Fadum *et al*. 2025), anammox will be completely excluded by 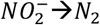 denitrifiers (Fig. S9). Therefore, in principle, under nitrite limitation, anammox bacteria completely exclude 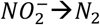 denitrifiers. In contrast, if anammox is limited by ammonium, both dependent groups can coexist. In this case, the 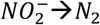 denitrifiers remain limited by nitrite. In fact, for the range of the supply of ammonium considered in our model, anammox is always limited by ammonium. Thus, anammox and 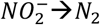 denitrifiers coexist as long as the nitrite production is sufficient for 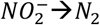 denitrifiers’ persistence at high supply of OM and nitrate (Fig. S8).

Because 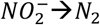 denitrifiers assist their feeder in suppressing the niche of 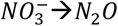 denitrifiers, whereas anammox exerts no such effect, the presence of anammox—which suppresses 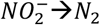 denitrifiers—indirectly benefits the feeder’s competitor ( 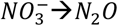denitrifiers), allowing them to expand their niche. Consequently, the coexistence space of 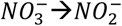 and 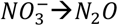 reducers broadens (Fig. 2acd). This, in turn, leads to an expanded N-loss region characterized by mixed *N*_2_ and *N*_2_*O* production.

#### Adding *N*_2_*O* → *N*_2_ denitrifiers: New Recipients Erode Feeder Dominance

When adding ***N***_**2**_ ***O* → *N***_**2**_ denitrifiers into the baseline system, the *N*_2_*O* **→** *N*_2_ denitrifiers depend on *N*_2_*O* produced by the 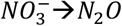 reducer. They also consume OM, but more efficiently than their feeder as indicated by their lower subsistence requirement 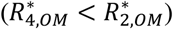. This is different from the previous subsystems, because the feeder, 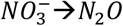 is a weaker competitor for OM when coexisting with the 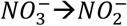 denitrifier. Consequently, introducing the *N*_2_*O* **→** *N*_2_ denitrifiers further increases the overall OM demand beyond that of the feeder alone. The consumption vector of this consortium is given by (Eqn. S94)

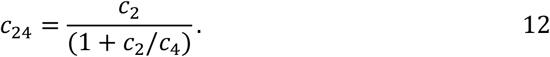

Because *c*_24_ is smaller than *c*_2_, the consortium has a stronger requirement for OM. This shift effectively reduces the competitive advantage of the feeder and expands the niche space of the 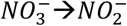 denitrifiers (Fig. 2e).

In biogeochemical terms, the *N*_2_*O* **→** *N*_2_ denitrifiers convert *N*_2_*O* to *N*_2_. In the virtual chemostat, *N*_2_*O* acts as an intermediate that is consumed by *N*_2_*O* **→** *N*_2_ denitrifiers immediately as it is produced. Thus, nitrogen loss is primarily in the form of *N*_2_ while still maintaining the potential for *N*_2_*O* escape (Fig. 3e).

### 3.3. The Full Ecological System and Its Biogeochemical impact Ecological Dynamics

The ecological result is a community structured by trade-offs, priority effects, and indirect facilitation, rather than stable coexistence of all possible functional types (Fig. 4a). When all seven functional groups are included, the system does not support coexistence of all types at steady state in the supply range of OM and nitrate considered in the model. Instead, in the regimes where most types (except B6 and B7) are likely sustained priority effects between functional consortiums determine the outcome. The 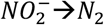 denitrifiers typically exclude the other recipient-feeder pair (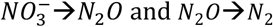 denitrifirs) by consuming OM more efficiently than the feeder 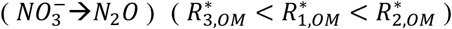, thereby restricting coexistence. However, anammox bacteria mitigated this dominance by outcompeting the 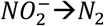 denitrifiers under nitrite limitation 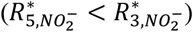, benefiting the recipient-feeder pair ( 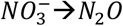 and *N*_2_*O*→*N*_2_ denitrifiers) and facilitating broader coexistence among denitrifiers. The extent of the benefit depends on the severity of ammonium limitation in anammox bacteria.

Complete denitrifiers (*B*_7_) only persist with sufficient supply of OM and are consistently outcompeted by more specialized groups when the supply of nitrate increases because of inefficient consumption on OM ( 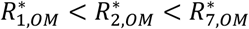, Table 1). For the supply range of OM considered in our simulation, it can only coexist with the 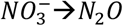 denitrifiers (*B*_2_).

The 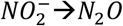 denitrifiers ( *B*_6_) could not exist when nitrite reducers or anammox are present under 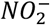 –limited condition. Because 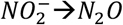 denitrifiers are a weaker competitor for nitrite compared to the nitrite reducers and anammox 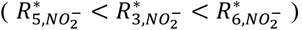. Thus, the 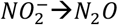 denitrifiers are effectively excluded, reflecting their weaker competitive ability.

#### Biogeochemical Impact

We next analyze the N loss patterns and other results relevant for the biogeochemistry of the “full system,” as described above, when all seven functional types are introduced into the virtual chemostat.

Sustaining any denitrifier population requires minimal supplies of both OM and nitrate 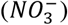; below these thresholds, no denitrifiers considered can persist in our model and nitrogen is conserved. Otherwise, the OM: 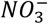 supply ratio strongly controls both the magnitude and form of nitrogen loss (Fig. 3b).

#### Net N_2_O Production Under Co-Limitation

Our findings identify a potential *N*_2_*O* accumulation regime. When both organic matter (OM) and nitrate 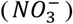 supplies are critically low, denitrifiers persist at densities too low to support downstream reducers, resulting in the production of *N*_2_*O* without subsequent consumption (Fig. 4b, lower-left corner). While previous studies suggest that increased OM availability generally promotes both *N*_2_*O* production and consumption (Li *et al*. 2022; Rummel *et al*. 2025), our results demonstrate that this relationship is also contingent upon 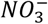 –availability. We show that when both electron donors and acceptors are scarce, the metabolic pools become jointly limiting, decoupling the production and consumption phases of the denitrification pathway. Under these conditions, the nitrogen flux proceeds only partially, accumulating *N*_2_*O* because the system lacks the reducing power or substrate flux necessary to sustain complete reduction to *N*_2_. Consequently, low substrate availability disrupts the metabolic balance between producers and reducers, fundamentally altering the magnitude and direction of the net nitrogen flux.

#### A Broad Niche for Anammox

A striking outcome of our model is the disparity in niche breadth between anammox bacteria and 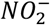 –reducing denitrifiers. Anammox occupy expansive nutrient supply regimes where 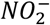 –reducers could persist, yet the reverse was not true. This asymmetry stems from our evidence-based assumption that anammox possesses a lower requirement for nitrite (a lower 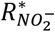) than its denitrifying competitors (but see (Fadum *et al*. 2025)). Consequently, if this competitive advantage holds, anammox maintains a strictly broader nutrient niche.

This ecological framework provides a mechanistic explanation for why anammox is more frequently detected in Oxygen Minimum Zones (OMZs), even in regions where denitrification-driven *N*_2_ production is prevalent (Dalsgaard *et al*. 2012; Babbin *et al*. 2014; Ward *et al*. 2019; Tracey *et al*. 2023). While denitrifiers are biogeochemically vital, their narrower niches likely render them less consistently detectable across varying environmental gradients.

More broadly, the two major *N*_2_ production pathways, 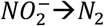 and anammox, have been observed to dominate different oceanic regions, leading to historical debate over their relative importance (Ward *et al*. 2009; Dalsgaard *et al*. 2012; Babbin *et al*. 2014; Tracey *et al*. 2023). Recent theoretical and modeling studies reconcile these contrasting observations by estimating that anammox contributes approximately 30% of global marine *N*_2_ production (Ward 2013; Babbin *et al*. 2014; Koeve *et al*. 2020; Zakem *et al*. 2020), with this fraction varying systematically in response to nutrient supply regimes (Sun *et al*. 2024).

## 4. Synthesis and General Principles

Our results reveal a set of general principles for how recipient populations reshape microbial coexistence and ecosystem function. Recipients may enhance the ecological niche of their feeders and/or their feeders’ competitor, or not, depending on the circumstances. The outcome depends on their competitive ability for limiting resources relative to both the feeder and its competitor as follows.

Reinforcement of feeder dominance occurs when recipients are specialized on resources that are not limiting to their feeders but limiting to the competitor of their feeders. In these cases, the feeder–recipient association intensifies competitive exclusion of the competitor (e.g., the group of *B*_1_, *B*_2_, and *B*_3_). In contrast, erosion of feeder dominance, by expanding feeder’s competitor niche, arises when recipients overlap strongly in resource use with their feeders but are more competitive for the limiting resource to their feeders. This reduces the feeder’s advantage and expands the niche of its competitor (e.g., the group of *B*_1_, *B*_2_, and *B*_4_). Finally, intuitively, no change in the dominance of the feeder occurs when recipients rely on resources unrelated to their feeder’s limitation, leaving competitive boundaries unaffected (e.g., the group of *B*_1_, *B*_2_, and anammox *B*_5_).

Together, these principles extend coexistence theory to multi-resource systems with dependencies. Results are relevant for complex microbial interaction networks beyond just that of the marine nitrogen cycle, providing a framework by which to untangle the complexities. Specifically, our results highlight that facilitation interacts with competition to determine both community assembly and ecosystem-scale biogeochemical function.

## 5. Conclusion

We show that under nitrite-limited, fully anoxic conditions, anammox bacteria occupy a broader nutrient niche than 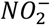 –reducing denitrifiers due to a lower 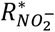. As a result, anammox can persist across all nutrient supply regimes that support denitrifiers, whereas the reverse is not true. This asymmetry provides a simple mechanistic explanation for the frequent detection of anammox in anoxic marine environments, including Oxygen Minimum Zones, even when denitrification-derived *N*_2_production is substantial.

Beyond niche structure, our results highlight how microbial interactions shape nitrogen loss pathways across nutrient regimes. Varying both OM and nitrate supply reveals richer coexistence patterns and biogeochemical outcomes than varying either resource alone, even when supply ratios are held constant. In this way, we extend (Sun *et al*. 2024) by demonstrating that increasing the OM: 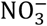 ratio drives transitions from complete denitrification to *N*_2_ toward partial denitrification across distinct nitrate supply regimes. Notably, under OM– and nitrate–colimited conditions, complete denitrification is constrained, allowing *N*_2_O to accumulate.

More broadly, our framework demonstrates how microbial traits and multi-resource supply jointly regulate ecosystem-scale nutrient transformations. Because recipient–feeder interactions are common across microbial systems, these insights extend beyond OMZs to diverse environments where coexistence governs the fate of energy and matter.

## References

1. Ali, M., Shaw, D.R., Albertsen, M. & Saikaly, P.E. (2020). Comparative Genome-Centric Analysis of Freshwater and Marine ANAMMOX Cultures Suggests Functional Redundancy in Nitrogen Removal Processes. Frontiers in Microbiology, Volume 11 – 2020.

2. Babbin, A.R., Keil, R.G., Devol, A.H. & Ward, B.B. (2014). Organic Matter Stoichiometry, Flux, and Oxygen Control Nitrogen Loss in the Ocean. Science, 344, 406–408.

3. Baeten, J.E., Batstone, D.J., Schraa, O.J., van Loosdrecht, M.C.M. & Volcke, E.I.P. (2019). Modelling anaerobic, aerobic and partial nitritation-anammox granular sludge reactors – A review. Water Research, 149, 322–341.

4. Bardgett, R.D. & van der Putten, W.H. (2014). Belowground biodiversity and ecosystem functioning. Nature, 515, 505–511.

5. Bertolet, Brittni L., Rodriguez, Luciana C., Murúa, José M., Favela, A. & Allison, Steven D. (2024). The Impact of Microbial Interactions on Ecosystem Function Intensifies Under Stress. Ecology Letters, 27, e14528.

6. Bruno, J.F., Stachowicz, J.J. & Bertness, M.D. (2003). Inclusion of facilitation into ecological theory. Trends in Ecology & Evolution, 18, 119–125.

7. Bulleri, F., Bruno, J.F., Silliman, B.R. & Stachowicz, J.J. (2016). Facilitation and the niche: implications for coexistence, range shifts and ecosystem functioning. Functional Ecology, 30, 70–78.

8. Chase, J.M. & Leibold, M.A. (2003). Ecological Niches: Linking Classical and Contemporary Approaches. University of Chicago Press.

9. Chesson, P. (2000). Mechanisms of Maintenance of Species Diversity. Annual Review of Ecology and Systematics, 31, 343–366.

10. Dalsgaard, T., Thamdrup, B., Farías, L. & Revsbech, N.P. (2012). Anammox and denitrification in the oxygen minimum zone of the eastern South Pacific. Limnology and Oceanography, 57, 1331–1346.

11. Fadum, J., Sun, X. & Zakem, E. (2025). Redox-constrained microbial ecology dictates nitrogen loss versus retention. ISME Communications, 5.

12. Falkowski, P.G., Fenchel, T. & Delong, E.F. (2008). The Microbial Engines That Drive Earth’s Biogeochemical Cycles. Science, 320, 1034–1039.

13. Fukami, T. (2015). Historical Contingency in Community Assembly: Integrating Niches, Species Pools, and Priority Effects. Annual Review of Ecology, Evolution, and Systematics, 46, 1–23.

14. Koeve, W., Kähler, P. & Oschlies, A. (2020). Does Export Production Measure Transient Changes of the Biological Carbon Pump’s Feedback to the Atmosphere Under Global Warming? Geophysical Research Letters, 47, e2020GL089928.

15. Lam, P. & Kuypers, M.M.M. (2011). Microbial Nitrogen Cycling Processes in Oxygen Minimum Zones. Annual Review of Marine Science, 3, 317–345.

16. Li, Y., Moinet, G.Y.K., Clough, T.J. & Whitehead, D. (2022). Organic matter contributions to nitrous oxide emissions following nitrate addition are not proportional to substrate-induced soil carbon priming. Science of The Total Environment, 851, 158274.

17. Orr, J.A., Armitage, D.W. & Letten, A.D. (2025). Coexistence Theory for Microbial Ecology, and Vice Versa. Environmental Microbiology, 27, e70072.

18. Rittmann, B.E. & McCarty, P.L. (2012). Environmental Biotechnology: Principles and Applications. Tata McGraw Hill Education Private Limited.

19. Rummel, P.S., Englert, P., Beule, L. & Pausch, J. (2025). N2O flux dynamics and production pathways modulated by soil organic matter and litter turnover. Biology and Fertility of Soils, 61, 1235–1251.

20. Sun, X., Buchanan, P.J., Zhang, I.H., San Roman, M., Babbin, A.R. & Zakem, E.J. (2024). Ecological dynamics explain modular denitrification in the ocean. Proceedings of the National Academy of Sciences, 121, e2417421121.

21. Sun, X. & Ward, B.B. (2021). Novel metagenome-assembled genomes involved in the nitrogen cycle from a Pacific oxygen minimum zone. ISME Communications, 1, 26.

22. Tilman, D. (1982). Resource competition and community structure. Monogr Popul Biol, 17, 1–296.

23. Tracey, J.C., Babbin, A.R., Wallace, E., Sun, X., DuRussel, K.L., Frey, C. et al. (2023). All about nitrite: exploring nitrite sources and sinks in the eastern tropical North Pacific oxygen minimum zone. Biogeosciences, 20, 2499–2523.

24. Ward, B.A., Collins, S., Dutkiewicz, S., Gibbs, S., Bown, P., Ridgwell, A. et al. (2019). Considering the Role of Adaptive Evolution in Models of the Ocean and Climate System. Journal of Advances in Modeling Earth Systems, 11, 3343–3361.

25. Ward, B.B. (2013). Oceans. How nitrogen is lost. Science, 341, 352–353.

26. Ward, B.B., Devol, A.H., Rich, J.J., Chang, B.X., Bulow, S.E., Naik, H. et al. (2009). Denitrification as the dominant nitrogen loss process in the Arabian Sea. Nature, 461, 78–81.

27. Xu, L., Klausmeier, C.A. & Zakem, E. (2025). Density dependence promotes species coexistence and provides a unifying explanation for distinct productivity-diversity relationships. bioRxiv, 2025.2007.2028.667229.

28. Zakem, E.J., Mahadevan, A., Lauderdale, J.M. & Follows, M.J. (2020). Stable aerobic and anaerobic coexistence in anoxic marine zones. ISME J, 14, 288–301.

29. Zhang, I.H., Sun, X., Jayakumar, A., Fortin, S.G., Ward, B.B. & Babbin, A.R. (2023). Partitioning of the denitrification pathway and other nitrite metabolisms within global oxygen deficient zones. ISME Communications, 3, 76.

